# Temporal irreversibility of neural dynamics as a signature of consciousness

**DOI:** 10.1101/2021.09.02.458802

**Authors:** Laura de la Fuente, Federico Zamberlan, Hernán Bocaccio, Morten Kringelbach, Gustavo Deco, Yonatan Sanz Perl, Enzo Tagliazucchi

## Abstract

Even though the fundamental laws of physics are the same when the direction of time is inverted, dissipative systems evolve in the preferred temporal direction indicated by the thermodynamic arrow of time. The fundamental nature of this temporal asymmetry led us to hypothesize its presence in the neural activity evoked by conscious perception of the physical world, and thus its covariance with the level of conscious awareness. Inspired by recent developments in stochastic thermodynamics, we implemented a data-driven and model-free deep learning framework to decode the temporal inversion of electrocorticography signals acquired from non-human primates. Brain activity time series recorded during conscious wakefulness could be distinguished from their inverted counterparts with high accuracy, both using frequency and phase information. However, classification accuracy was reduced for data acquired during deep sleep and under ketamine-induced anesthesia; moreover, the predictions obtained from multiple independent neural networks were less consistent for sleep and anesthesia than for conscious wakefulness. Finally, the analysis of feature importance scores highlighted transitions between slow (≈20 Hz) and fast frequencies (> 40 Hz) as the main contributors to the temporal asymmetry observed during conscious wakefulness. Our results show that a preferred temporal direction is simultaneously manifest in the neural activity evoked by conscious mentation and in the phenomenology of the passage of time, establishing common ground to tackle the relationship between brain and subjective experience.

## Introduction

Our everyday conscious experience invariably includes the perception of time continuously flowing from past to future, but never in the opposite direction (Kent and Wittmann, 2021). In physics, this asymmetry is indicated by the thermodynamic arrow of time, which determines a preferred direction for the temporal evolution of certain macroscopic systems, i.e. towards states of increased entropy (Eddington, 1928; Parrondo et al., 2009). While this is not true for all physical processes (for instance, the laws of physics are compatible with planets reversing their direction of movement around the Sun), it is valid for the majority of dissipative non-equilibrium phenomena that sustain life (Prigogine, 1975). How does this asymmetry of the physical environment translate into the temporal properties of its neural representation? Is the temporal irreversibility of neural dynamics related to the cognitive experience of the flow of time, and thus can it be altered in different states of consciousness?

The possibility of temporally reversible neural dynamics across conscious and/or behavioral states can be discarded from the outset, since brain activity evoked by sensory perception of the physical environment will naturally present a certain degree of temporal asymmetry. Indeed, emerging evidence supports the view that brain dynamics unfold outside of thermodynamic equilibrium, a scenario where the transition probabilities between states in configuration space are asymmetric, the entropy production rate is strictly positive, and neural activity time series are temporally irreversible (Lynn et al., 2020; Sanz Perl et al., 2021; Deco et al., 2021). Moreover, the degree of proximity to equilibrium dynamics carries functional relevance, since it is related to the level of consciousness (Sanz Perl et al., 2021), has been shown to change during the performance of effortful cognitive tasks (Lynn et al., 2020; Deco et al., 2021), and is altered in neuropsychiatric patients (Zanin et al., 2020).

After the possibility of reversible dynamics is discarded, we can distinguish at least three non-mutually exclusive answers to the question posed in the first paragraph. First, the temporal asymmetry of evoked neural responses could mirror the asymmetry present in consciously perceived sensory information. The use of naturalistic stimuli in human neuroimaging studies has established that brain activity time series measured with different modalities can show consistent correlations with features extracted from the stream of incoming sensory information, adding support to this possibility (Lankinen et al., 2018). Conversely, in the absence of conscious information processing, the responses evoked by sensory stimulation tend to be regular and stereotypical, lacking the temporal complexity indicative of irreversible dynamics (Czisch et al., 2004; Krom et al., 2020; Górska, et al., 2021). A second possibility is that macroscale brain activity is inherently asymmetric, regardless of how sensory inputs are structured in time. This is likely the case for activity recorded using functional magnetic resonance imaging (fMRI), since irreversible physiological processes (i.e. the hemodynamic response) mediate between the neural response and the recorded signal (Logothetis and Wandell, 2004). Finally, as brain development occurs while embedded within a temporally asymmetric environment, the emerging intrinsic activity patterns could present this asymmetry even during unconscious states (Pan and Monje, 2020). In this case, irreversibility could reflect a property of a dynamic baseline (Raichle, 2006), with brain activity unfolding in the proximity of functional patterns that facilitate rapid reactivity towards external demands (Deco et al., 2013), or as a consequence of neural activity replay necessary for learning and memory consolidation (Deuker et al., 2013).

To disentangle these extrinsic and intrinsic potential sources of temporal irreversibility, we investigated electrocorticography (ECoG) time series obtained from non-human primates during conscious wakefulness and two states of reduced consciousness: deep sleep and ketamine-induced general anesthesia. We note that reduced or absent temporal asymmetry of ECoG time series during unconscious brain states would be compatible with the first scenario outlined in the previous paragraph, while preserved temporal asymmetry during unconsciousness would be compatible with the second and third possibilities. We assessed the temporal reversibility of ECoG signals adapting a model-free machine learning framework from stochastic thermodynamics, where a deep convolutional network was trained and tested to distinguish short segments of signals from their temporally inverted counterparts (Seif et al., 2021).

## Results

The outline of the procedure for ECoG dimensionality reduction and estimation of temporal reversibility is shown in Fig. 1, including sample electrode locations (Fig. 1A), percentage variance explained vs. number of principal components (Fig. 1B), an illustration of forward and inverted time series (Fig. 1C), and of the different inputs for the deep convolutional network (Fig. 1C) whose architecture is represented in Fig. 1E. We trained and tested a total of 3×3×3×3= 81 different deep learning models for classification of forward vs. inverted ECoG time series. There are three possible choices for the number of principal components used for analysis (1st, 1st+2nd, and 1st+2nd+3rd principal components); another three choices for neural network complexity (i.e. number of convolutional blocks), another for the data type used as input to the neural network (frequency, phase, or both combined), and finally three states of consciousness (wakefulness, deep sleep and ketamine anesthesia). Each model was trained to distinguish between forward and inverted versions of short segments of ECoG data (Seif et al., 2021; Deco et al., 2021). The area under the receiver operating characteristic curve (AUC) and the loss in the validation sample were calculated for all models as metrics of classifier performance. Null models were constructed to determine if the loss was significantly larger than expected at chance level, i.e. whether the loss of the classifier without label shuffling was consistently better than the one obtained for classifiers trained with label shuffling.

**Figure 1.**
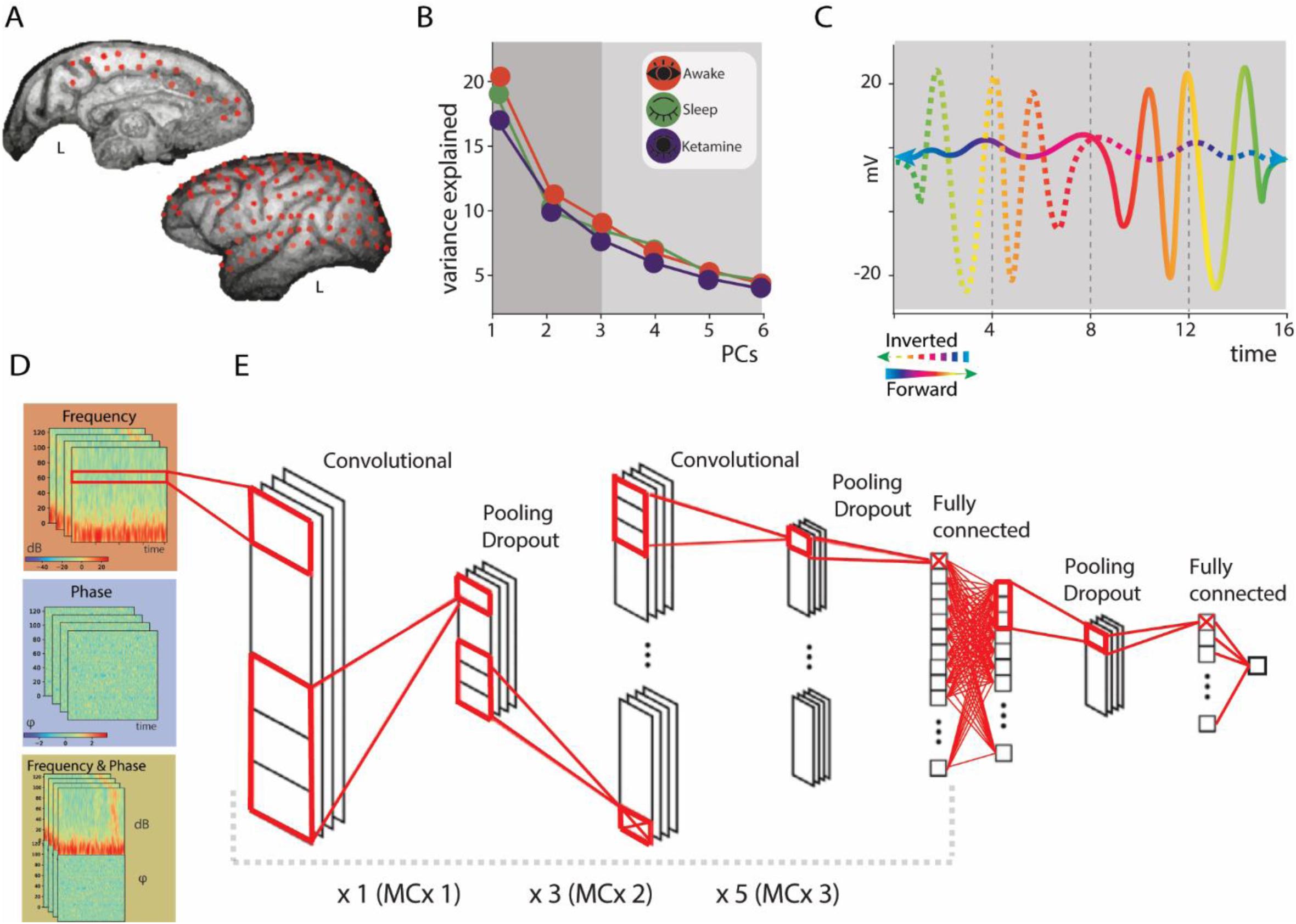
Procedure for dimensionality reduction and estimation of temporal reversibility. **A**. Sample locations of the ECoG electrodes. **B**. Percentage of variance explained as a function of the number of principal components for wakefulness, sleep and ketamine anesthesia. **C**. Illustration of the forward (solid) and inverted (dashed) versions of a signal. **D**. Examples of data used as input to the deep learning classifier: frequency (red), phase (blue), and both combined (yellow). Each time series was entered as an independent channel of a 1D deep convolutional network. **E**. Illustration of the neural network architecture used to classify forward vs. backward signals. Layers enclosed within a grey dashed line were repeated 1, 3 and 5 times for models of complexity 1, 2 and 3, respectively (here termed MCx 1, MCx 2 and MCx 3).

Figure 2 shows the results obtained for all deep learning models. Forward vs. inverted wakefulness ECoG signal segments could be distinguished with performance significantly above chance for all deep learning models with AUC > 0.75. However, this did not occur for sleep data with frequency as input (significant performance was obtained only for the MCx 3 model, with 1st+2nd components as input). For ketamine anesthesia, all the models of low complexity based on frequency data failed to yield performance significantly above chance level, but higher complexity models performed similar to those trained with wakefulness data. Models trained using the phase of ketamine ECoG data systematically failed to perform above chance, while those trained using the phase of sleep ECoG data performed well only if all three first principal components were included. The same trend of decreased performance vs. number of principal components and model complexity was observed for models trained with frequency and phase information combined. In summary, while all models performed very well when classifying forward vs. inverted wakefulness ECoG data, their performance was much less consistent for sleep and ketamine, resulting in chance level accuracy except for certain high complexity models.

**Figure 2.**
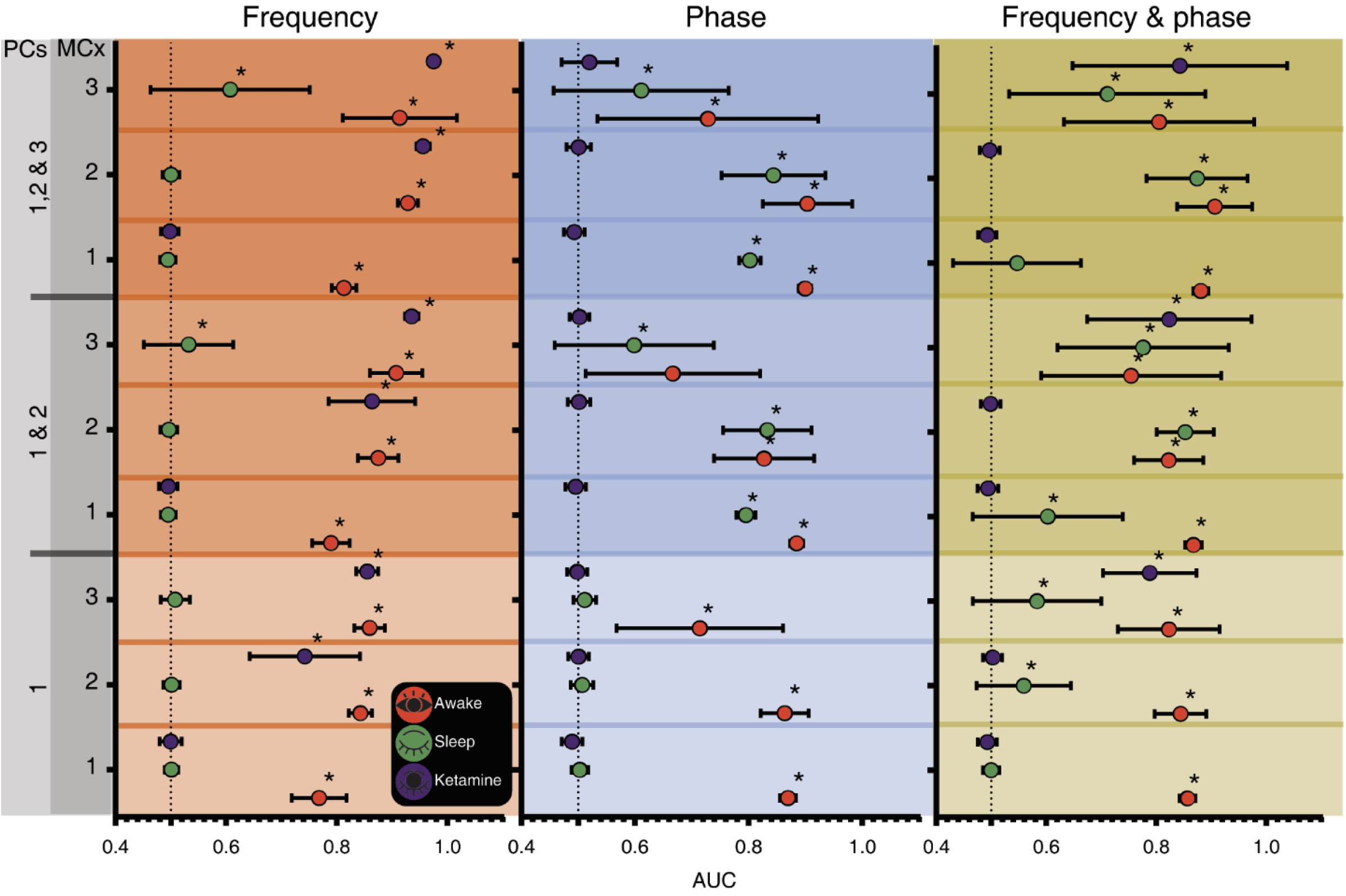
Classifier performances depend on the state of consciousness. Each point (mean ± standard deviation) indicates the AUC of one of the 81 deep learning models trained to distinguish forward vs. inverted signals. Columns indicate the data type used as input (frequency, phase, and both), while rows are ordered based on the number of principal components used for analysis (PCs), and for each selection of PCs, according to the model complexity (MCx). Asterisks indicate that the loss without label shuffling was less than the loss obtained with label shuffling across all 1000 iterations (p<0.001).

Next, we assessed how the consistency of the model predictions depended on the state of consciousness. We obtained the output of the last neural network layer for all ECoG epochs of each state, which is interpreted as the probability of that epoch being temporally inverted. We then obtained the correlation between these probabilities for different neural network architectures. Figure 3 shows these correlations for all states of consciousness (rows) and input data types (columns). All correlation values were close to 1 for conscious wakefulness, and significantly higher than those obtained for sleep and ketamine-induced anesthesia (as determined using Fisher’s z and Zou’s confidence intervals implemented in the R cocor package, see Methods). This result implies that time reversal during wakefulness can be detected with higher consistency relative to sleep and ketamine anesthesia.

**Figure 3.**
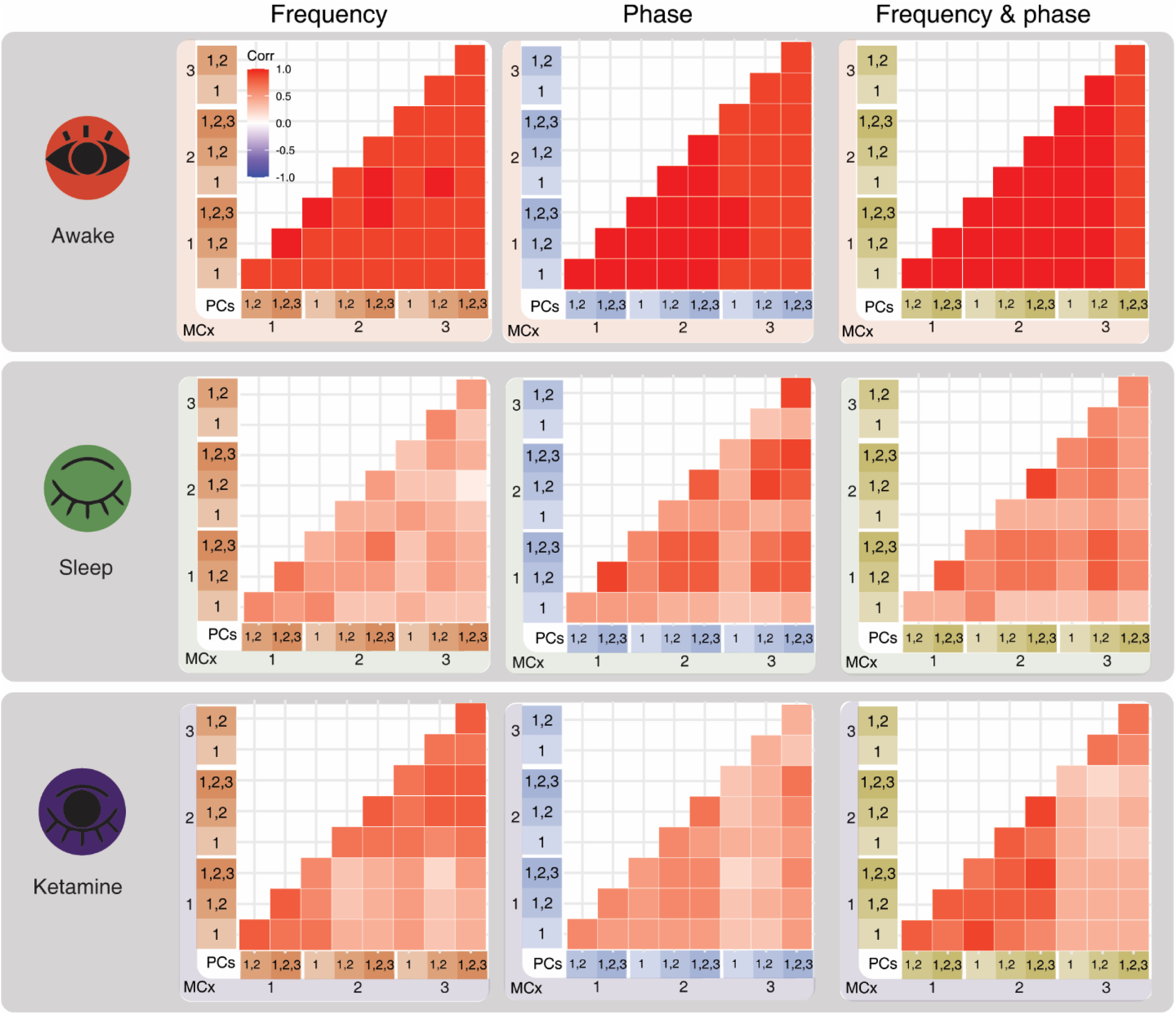
The consistency between models depends on the state of consciousness. Spearman correlations (p < 0.05, FDR corrected) between the predictions obtained from models with different number of principal components (PCs) and complexity (MCx), for all states of consciousness (rows) and input data type (columns). All correlations for wakefulness were significantly larger than those obtained for sleep and ketamine-induced anesthesia.

**Figure 4.**
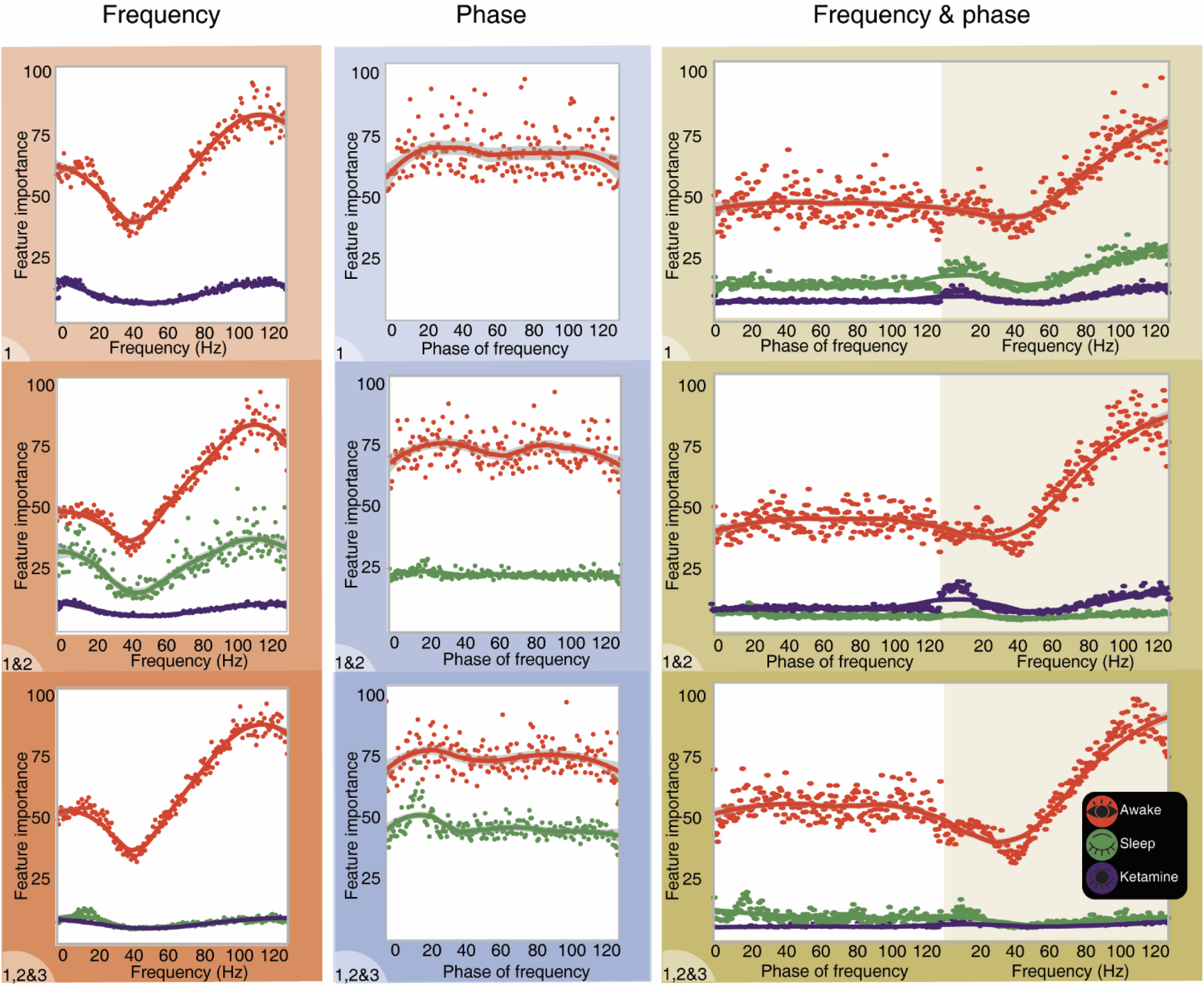
Feature importance scores vs. frequency. Average absolute value of the integrated gradients for each data channel comprising frequency (left, red), phase (center, blue) and both (right, yellow), and for each conscious state (awake, sleep and ketamine-induced anesthesia). The integrated gradients were computed for samples correctly classified in at least 75% of the models, and only for models that performed better than their null counterparts constructed by label shuffling. In all cases, the highest relative contribution was obtained for high (>40 Hz) frequency bands with a secondary peak in the beta band (≈20 Hz), while the feature importance corresponding to phase information did not present clearly discernible variations.

Finally, we investigated the characteristics of the time series that were most relevant for the detection of time reversal. We identified samples that were correctly classified with AUC significantly larger than 0.5 (assessed via label shuffling) in at least 75% of the deep learning models. We then computed the integrated gradients (a feature importance metric; Sundararajan et al., 2017) and averaged their absolute values for each data channel and state of consciousness. We found that models trained using wakefulness data presented higher values, both for frequency and phase. In the frequency domain, a greater relative importance was observed for high (>40 Hz) frequencies within the gamma band, with a secondary peak in the beta band (≈20 Hz). This was not observed for models trained using phase information, where all features carried similar importance. These results were maintained for the models trained using combined frequency and phase information.

## Discussion

A sense of being situated both in time and space is a fundamental aspect of conscious experience, and arguably a prerequisite for anticipating and planning for the future. Throughout their evolutionary history, brains have acquired the capacity to construct multiple and increasingly sophisticated representations of the physical environment (Gross and Graziano, 1995; Kragel et al., 2018), which interact with mental models of the world to predict the consequence of motor actions (Gilead et al., 2020). The temporal properties of physical events (such as their duration) are potentially unique, in the sense that their neural representation can be based on the very property that is being represented (Northoff and Huang, 2017; Northoff et al., 2020). Thus, while temporal intervals between external events can be mapped to temporal intervals between neural events, properties such as color or scent are not mapped to neurons that are colored or have specific aromas (note that the general validity of this assumption has been contested, see Dennett and Kinsbourne, 1992). This intimate relationship between time and its neural representation suggests that the temporal asymmetries present in the environment of an organism should leave an imprint in its brain activity dynamics. We investigated the nature of this imprint, distinguishing between different possible scenarios, and assessing their likelihood by analyzing neural dynamics recorded during conscious and unconscious states.

In natural environments, conscious perception almost certainly includes processes that are temporally irreversible, such as the majority of meaningful sounds used for orientation or communication. Information that is consciously perceived is capable of exerting a global influence on the spatial and temporal organization of brain activity (Mashour et al., 2020); thus, it can be expected that the ongoing processing of conscious sensory percepts breaks the temporal symmetry of large-scale neural dynamics. Conversely, when the cortex becomes isolated from the environment, self-generated activity patterns are regular and appear to be reversible in time, as is the case of the high amplitude slow oscillations that emerge during deep sleep or under the effects of certain anesthetic drugs (Sanchez-Vives and Mattia, 2014; Timofeev et al., 2020). While intuitively appealing, the potential link between temporal asymmetry and conscious awareness has remained largely unexplored: few studies to date addressed the reversibility of brain activity time series (Zanin et al., 2020; Deco et al., 2021), and none attempted to link this reversibility to the global state of consciousness.

We adopted a data-driven and model-free approach to assess the breaking of temporal symmetry in ECoG time series obtained from non-human primates undergoing different states of consciousness. This approach is based on training deep convolutional networks to distinguish samples with and without temporal inversion, and then estimating their out-of-sample generalization capacity. A similar procedure was adopted in recent articles showing that machine learning models are capable of representing the direction of the flow of time in macroscopic (Wei et al., 2018) and microscopic processes (Seif et al., 2021). As expected, the capacity of the network to detect temporal inversion increased with its complexity (i.e. number of convolutional blocks) and with the amount of information provided as input (from one to three principal components of the ECoG data). Nevertheless, all AUC values were indicative of chance level classification accuracy for ketamine anesthesia (using phase data as input) and for deep sleep (using frequency data as input). Only the temporally-inverted epochs of wakefulness ECoG data could be reliable detected by all neural networks, regardless of their complexity and amount of input data.

We showed that ECoG activity present in fast (>40 Hz) frequencies was the main contributor to temporal irreversibility during wakefulness. This frequency range corresponds to the gamma band, which has been implicated in several aspects of sensory and cognitive processing in both human and non-human primates (Başar-Eroglu et al., 1996; Jia and Kohn, 2011). Importantly, oscillations in the gamma band have been proposed as fundamental for the flexible and dynamic inter-areal routing of information in the brain (Fries, 2015; Palmigiano et al., 2017). As sensory data is processed by a hierarchy of increasingly specialized cortical regions, the gamma band plays an important role in the binding of low-level information into moving objects and dynamic scenes (Singer, 2001). Thus, if the stream of incoming sensory data evolves in a preferred temporal direction, it can be expected that this preference will also manifest in the dynamics of gamma band oscillations. In contrast, internally generated oscillations (typically involving interactions with subcortical structures, such as the thalamus and the reticular formation) are more regular and lack the temporal complexity characteristic indicative of irreversible processes (Sanchez-Vives and Mattia, 2014; Timofeev et al., 2020).

It must be noted that some of the trained models were successful to identify time reversal in presumably unconscious brain states, such as deep sleep and ketamine-induced anesthesia. There are multiple (possibly overlapping) explanations for this result. First, even though conscious perception is absent in these two states, conscious mentation might nevertheless be present in the form of vivid “dreams” (Siclari et al., 2017; Ballesteros et al., 2020). Thus, if temporal irreversibility is a characteristic of conscious information processing, then it could be expected in these two states, at least sporadically. Second, intrinsic brain activity might recapitulate functionally meaningful patterns even in the absence of conscious awareness (Sadaghiani et al., 2010). The elevated metabolic cost of this activity is consistent with the important role of continuously constructing and testing generative models of the world (Koren and Denève, 2017); in this case, the temporal asymmetry of neural dynamics could reflect the structure of those models. Finally, phase information was useful to detect temporally inverted epochs during sleep, which could reflect temporal asymmetries caused by slow travelling oscillations with well-defined sinks and sources (Murphy et al., 2009).

There is a close relationship between temporal reversibility and thermodynamic equilibrium. The latter does not refer to equilibrium in the mechanical sense (i.e. absence of accelerated movement), but to the presence of stochastic fluctuations that are compensated over time, requiring that the likelihood of observing a certain trajectory in configuration space is equal to the likelihood of observing its opposite (Seifert, 2008). In living systems, departures from non-equilibrium dynamics are scale-dependent; for example, the intracellular milieu might be at equilibrium, while at the same time the flagella continuously undergo an irreversible cycle which propels the cell (Gnesotto et al., 2018). The same observation applies to neural processes and thus to the temporal asymmetry of brain activity time series. The dynamics of single neurons are irreversible, since temporal symmetry is broken by the process underlying the generation of action potentials. However, this symmetry might be recovered at coarser scales, where the activities of single cells are averaged to result in oscillatory local field potentials (Sanz Perl., 2021). As previously discussed, some of these oscillations (e.g. the gamma rhythm) can in turn be modulated to represent information associated with temporally irreversible physical processes. Also, while ECoG recordings provide direct measurements of neuroelectric activity, less direct neuroimaging modalities might present temporal asymmetries which do not originate from neural sources, as is the case of the hemodynamic response in fMRI (Logothetis and Wandell, 2004).

Our results can be seen as the first steps towards a framework to disentangle the intrinsic and extrinsic factors that contribute to spontaneous and evoked neural dynamics, and how each of them pushes the brain towards and away from equilibrium. Animal experiments with cortical deafferentation (i.e. removal of thalamocortical inputs) show that macroscale neuroelectric activity drifts towards stable and regular dynamics in the absence of extrinsic stimulation (Lemieux et al., 2014; Timofeev et al., 2020). On the other hand, the critical brain hypothesis postulates that complex endogenous dynamics do not necessitate continuous external forcing (Cocchi et al., 2017; Chialvo, 2010), although the evidence in favor of slightly sub-critical dynamics suggests that a certain measure of extrinsic activity is required to maintain the brain away from equilibrium (Priesemann, et al., 2014). Future work should simultaneously investigate the temporal symmetry of ongoing and evoked brain activity, together with that of the chosen sensory stimuli, while conducting this analysis for different global brain states characterized by their natural tendency to move towards or away from equilibrium. Understanding how temporal asymmetries propagate from stimuli to evoked activity in the presence of endogenous fluctuations could be especially relevant for future clinical developments, since some of the most devastating neurological conditions are associated with abnormal shifts towards more stable dynamics (Casali et al., 2013).

In summary, we adapted a deep learning framework previously used in stochastic thermodynamics to show that the irreversibility of large-scale brain dynamics changes from wakefulness to unconscious states. Due to their uniquely straightforward neural representations, temporal properties might constitute valuable common ground between physics and phenomenology, with the potential to open new directions of research in the neuroscientific study of consciousness.

## Materials and methods

The data and code used in this manuscript are feely available at https://github.com/Lauralethia/TENET.git.

### Data

We based our analysis on electrocorticography (ECoG) recordings obtained from non-human primates in different states of consciousness (awake, 2 animals, 8 sessions; deep sleep, 2 animals, 21 sessions and under anesthetic doses of ketamine,∼5.0 mg/kg, 2 animals, 4 sessions) (Yanagawa et al., 2013). Recordings were obtained from 128 channels, with an example layout shown in Fig. 1A. All the analyzed sessions were conducted with eyes closed. For more information on the dataset visit http://neurotycho.org/.

### ECoG data preprocessing

The first preprocessing step consisted of a notch filter to eliminate line noise and its harmonics (50Hz, 100Hz, 150Hz); next, time series were bandpass filtered between 5 and 500Hz, and resampled from 1 KHz to 256Hz. Following previous work (Sanz Perl et al., 2021), principal component analysis was applied to reduce the dimensionality of each ECoG recording, retaining up to the first three principal components which cumulatively explained ≈40% of the variance (Fig. 1B). These time series were divided into 4 s epochs, resulting in a total of 3759 epochs for wakefulness, 4198 epochs for deep sleep, and 2556 epochs for ketamine. Each epoch was then inverted in time, giving a forward and an inverted sample per epoch.

### Feature construction and selection

The time series corresponding to the three principal components were converted to frequency and phase representations using a fast Fourier transform with a window of 32 overlapping time points and a hop length of 4 points in 201 intervals, covering the frequency range up to 128 Hz. The frequency data was transformed to decibels. This information, computed using the forward and backward signals, comprised all the features associated with each individual sample. However, different feature subsets were used to train and test the deep learning classifiers. These subsets were given by all combinations of frequency, phase, and frequency plus phase, obtained for the 1st, 1st and 2nd, and 1st, 2nd and 3rd principal components (Fig. 1D). For example, a possible classification problem could consist of distinguishing between forward and backward wakefulness epochs based on the phase information of the 1st and 2nd principal components.

### Neural network architecture

We trained 1D convolutional neural networks (implemented in PyTorch, https://pytorch.org/) to distinguish between forward and inverted epochs based on the information described in the previous subsections. Each input channel corresponded to the frequency and or/phase time series, with 201 channels per principal component; for instance, a model trained using frequency and phase information of the first two components would have a total of 804 input channels (201 frequency channels and 201 phase channels per principal component).

The architecture of the deep learning models is illustrated in Fig. 1E. In all models, the first convolutional layer dropped the number of input channels to 100, and the last layer consisted of a single unit with a sigmoid activation function. We constructed models with three levels of complexity (MCx 1, 2 and 3), which included 1, 3 and 5 convolutional blocks, respectively. Each convolutional block consisted of batch normalization, followed by a 1D convolutional layer with 4-point filters, another batch normalization, 1D max pooling (kernel size = 2, stride = 2, without padding) and a dropout layer (p = 0.4). After the convolutional block, a fully connected layer reduced the number of features (12700 for MCx 1, 12000 for MCx 2 and 9600 for MCx3) to 1024. A final dropout layer (p=0.1) was followed by a dense layer ending in a single sigmoid neuron trained using binary cross entropy loss with classes 1 (forward) and 0 (inverted). Biases were not optimized and exponential linear units (ELU) were used as activation functions. The details of the neural network implementation can be found in the repository (https://github.com/Lauralethia/TENET.git)

### Deep learning models

A total of 81 different models were constructed, resulting in the combination of 3 model complexity levels (MCx 1 to 3), using 3 types of data (frequency, phase, and both) computed from the 1st, 1st and 2nd, and 1st, 2nd and 3rd principal component time series. Each model was repeated for 100 iterations, with and without label shuffling to construct a null model. During training, the data were divided into five folds, one used for validation and the others for training with a batch size of 1800 examples in 200 epochs using and an exponential cyclical learning rate (basal 5e-5, maximum 5e-4), 100 increasing and 100 decreasing with an Adam optimizer with weight decay of 1e-3. In each iteration, the loss, accuracy, and area under the curve (AUC) were recorded to determine model performance both with and without label randomization. To summarize the training after the 100 iterations of each model, the model with the lowest loss was selected and the AUC for randomized vs. non-randomized labels was compared with a one-side Welch t-test, being considered significant at p < 0.001.

### Consistency between models

The model consistency was evaluated by computing the Spearman correlation coefficient between the output probabilities obtained for models of different complexity and number of principal components. Correlations were considered significant at p < 0.05, False Discovery Rate (FDR) corrected. To compare consistency (i.e. correlations) between models, we used the *cocor* R package (http://comparingcorrelations.org/), which implements Fisher’s z and Zou’s confidence intervals to estimate statistically significant differences between correlation coefficients of independent samples given their corresponding sample sizes (Diedenhofen and Musch, 2015).

### Feature importance

We selected a subset of samples that were correctly classified by 75% or more of the models with the purpose of investigating which features were the most relevant for the classification. We calculated the integrated gradients (Sundararajan et al., 2017) and then computed the average absolute value for each feature (frequency, phase, or frequency and phase) for each conscious state, interpreting this result as the feature importance for the binary classification problem.

## Acknowledgements

This work was supported by funding from Agencia Nacional De Promocion Cientifica Y Tecnologica grants PICT-2018-03103 and PICT-2019-02294. L.F., F.Z. and H.B. were supported by a doctoral fellowship granted by the National Scientific and Technical Research Council (CONICET). E.T. was supported by a Mercator Fellowship awarded by the DFG. The authors acknowledge Christopher Nolan CBE for indirectly inspiring the core idea of this manuscript.

